# How Phosphorylation of α/β-Tubulin Perturbs Microtubule Structure: A Computational Study

**DOI:** 10.64898/2026.04.29.721677

**Authors:** Annemarie Ianos, Ahmed Osman, Baofu Qiao, Susan A. Rotenberg

## Abstract

Microtubules are cytoskeletal structures composed of polymers of α/β-tubulin heterodimers that enable cell division and motility by a process of alternating episodes of polymerization and depolymerization (dynamic instability). Transition from a polymerizing to a depolymerizing microtubule is triggered at the interdimer interface by Glu254 in α-tubulin (α:Glu254), which hydrolyzes GTP bound to β-tubulin (β:GTP). The process is regulated by phosphorylation of α-tubulin (Ser165) or β-tubulin (Ser172) via signaling protein kinases (PKC, CDK1). All-atom molecular dynamics simulations of α/β-tubulin 6-mer systems are used to screen the cryo-EM structure of a microtubule (PDB 3J6E) for structural responses to phosphorylation of each tubulin subunit. In terms of global structure, microtubules with phosphorylated α-tubulin have a straight conformation attributed to a growing microtubule, whereas MTs with phosphorylated β-tubulin are curved, characteristic of a disassembling MT. Phospho-α-tubulin initiates displacement of key secondary structures (helix H8, loop T5) at the inter-dimer interface, shifts the β:GTP nucleotide by 5 Å, and immobilizes the γ-phosphate of β:GTP through increased H-bonding with β-tubulin. Phospho-β-tubulin produces fewer structural effects and has a more flexible β:GTP. For β:GTP hydrolysis, the phospho-β-tubulin system displays an extensive network of water molecules between α:Glu254 and the γ-phosphate of β:GTP, facilitating its hydrolysis. In contrast, phospho-α-tubulin displays a discontinuous network of water molecules that predicts a diminished capacity for β:GTP hydrolysis. These findings provide a detailed framework for understanding how phosphorylation of each tubulin subunit restructures the inter-dimer interface to modulate β:GTP hydrolysis, global structure, and dynamic instability in response to key signaling protein kinases.

## INTRODUCTION

Microtubules (MTs) are components of the cytoskeleton that mediate essential cell functions such as cell movement and cell division (1). Mammalian MTs are composed of α- and β-tubulin heterodimers arranged through longitudinal and lateral interactions to form the MT as a hollow cylindrical lattice (**Figure 1**) (2). MTs exhibit a stochastic process of alternating polymerization and depolymerization episodes referred to as “dynamic instability” (3, 4). Elongation occurs by adding α/β-tubulin heterodimers to the growing ends of protofilaments (“plus ends”) and creating a β-tubulin-bound guanosine triphosphate (β:GTP) “cap”.

**Figure 1.**
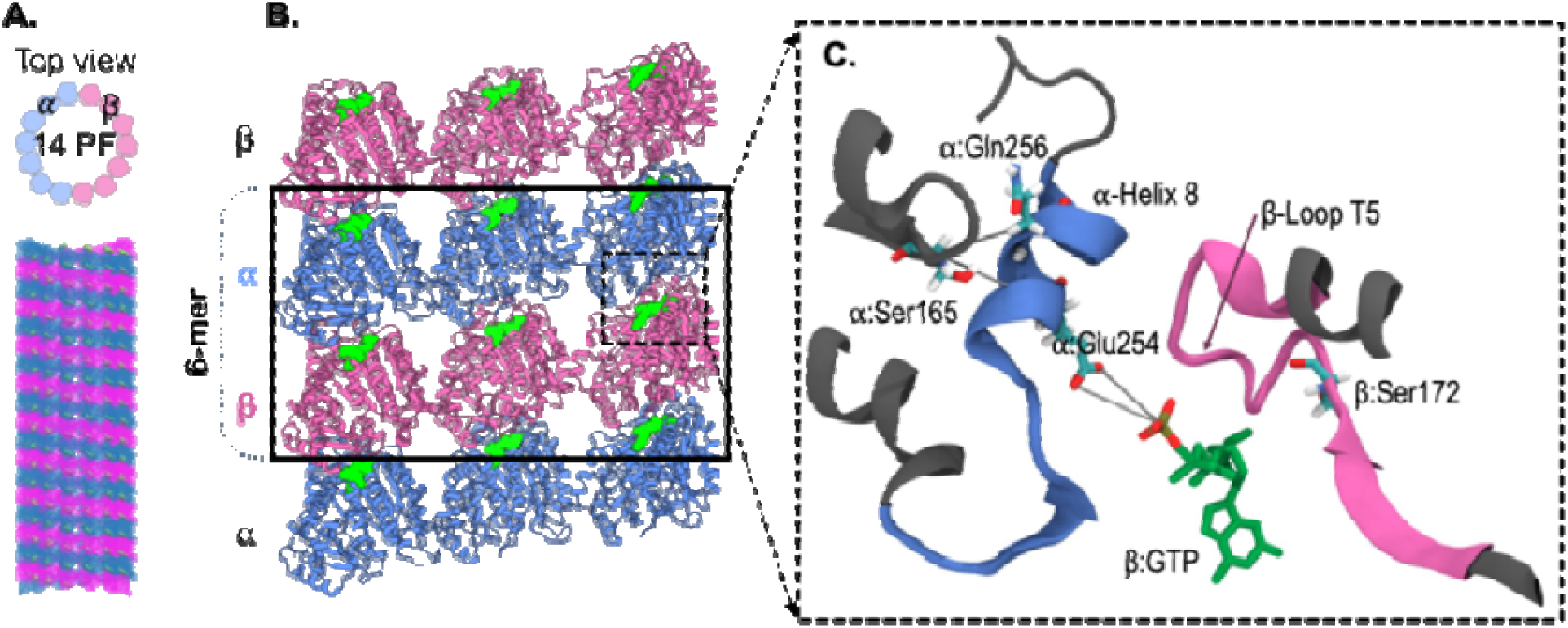
Structure of microtubules and the 6-mer model used in simulations. A) Scheme depicting an MT consisting of 14 protofilaments (PFs). B) The 6-mer system consisting of fragments of three protofilaments of an infinitely long MT. C) The heterodimer interface containing helix H8 and loop T7 of α-tubulin (blue) and loop T5 (magenta) of β-tubulin and critical residues. These secondary structures are positioned near the γ-phosphate of the E-site β:GTP.

Hydrolysis of GTP to GDP causes MTs to undergo rapid and sudden disassembly (“catastrophe”), whereas replacement with a new GTP molecule resumes polymerization (“rescue”) (5, 6). During these transitions, hydrolysis of the exchangeable β:GTP to GDP occurs at the interdimer cleft. In contrast, α-tubulin-bound GTP, which is located on the intradimer, non-exchangeable N-site, is not hydrolyzed (6). A key player in the hydrolysis of β:GTP is α:Glu254, the sole catalytic residue in α-tubulin located at the inter-dimer cleft (**Figure 1**). The essential catalytic role of α:Glu254 was demonstrated with an α:E254A mutation that gave rise to persistent MT elongation (7). Thus, the catalytic availability of α:Glu254 to perform hydrolysis of β:GTP contributes to the regulation of MT dynamic instability.

In an earlier study from this laboratory (8), α-tubulin was discovered to be phosphorylated by protein kinase C (PKC) at Ser165 (α:Ser165) in non-transformed human breast cells (MCF-10A cells). Specific phosphorylation of α-tubulin by a PKC activator (8) correlated with increased cell motility, which was earlier found to be coincident with suppressed proliferation (9). Measurement of MT dynamics in these cells demonstrated that phosphorylation of α:Ser165 promotes enhanced rates (by 1.5-fold) and duration (by 3-fold) of the growth phase with decreased occurrence of catastrophe (10). These parameters were reproduced with a phospho-mimetic mutation at Ser165 in α-tubulin (α:S165D) and suppressed by a mutation that blocked phosphorylation (α:S165N).

Since MCF-10A cells express low levels of phospho-α-tubulin (due to low PKCα expression), the impact of engineered over-expression of α:S165N-tubulin was explored in metastatic breast cancer cells (MB-MDA-231). These cells express moderate levels of phosphorylated α-tubulin and respond to the competitive effect of α:S165N-tubulin with suppressed motility (9). Unexpectedly, these cells produced hyperproliferative tumors in mice (11). These findings led us to consider a ‘toggle-switch’ model for describing how phosphorylation/dephosphorylation of α:Ser165 modulates MT dynamics and consequently cellular phenotypes.

Owing to the mechanistic connection between the state of phosphorylation of α:Ser165 and MT dynamics (9), we explored whether phosphorylation of α-tubulin perturbs access to β:GTP by the catalytic α:Glu254 residue, thereby resulting in a lower rate of hydrolysis. Inspection of the molecular structure of an α/β-tubulin polymer revealed that α:Ser165 and the binding site for β:GTP occur within 11 Å of each other near the longitudinal interface between two consecutive α/β-tubulin heterodimers (inter-dimer cleft) (12). Positioned near this interface in α-tubulin is helix H8, which contains catalytic α:Glu254 whose carboxylates are located 6-7 Å from the γ-phosphate of β:GTP. An attractive hypothesis is that phosphorylation of α:Ser165 leads to an altered spatial relationship between α:Glu254 and β:GTP, consequently impairing β:GTP hydrolysis; this impediment would prolong MT elongation and thus explain the observed MT dynamics (9) and increased cell motility observed with α-tubulin phosphorylation (9). This idea was recently addressed with simulations using a cryo-EM structure of an MT to model the effects of α:S165D compared with that of α:S165N (13). Findings with the α:S165D system showed that the distance between α:Glu254 and the β:GTP γ-phosphate increased by 0.6 Å while the β:GTP nucleotide base was rotated by 4 Å. Furthermore, in immunocytochemistry experiments with MCF-10A cells that had been either genetically engineered to express α:S165D or treated with a PKC activator, an antibody recognizing GTP-bound tubulin demonstrated significantly enhanced signals in both conditions (13). These findings are consistent with a disruption in the spatial relationship between α:Glu254 and β:GTP that in turn impedes β:GTP hydrolysis.

In the present study, we extend the previous computational analysis by incorporating the effects of β-tubulin phosphorylation, an event mediated by cell cycle-dependent kinase-1 CDK1 (14). This enzyme was previously shown to increase cell proliferation in ovarian cancer cells through its phosphorylation of β-tubulin at β:Ser172 (14). The reciprocal effects on proliferation by phospho-α-tubulin and phospho-β-tubulin prompt us to expand the toggle switch model to incorporate alternating states of phosphorylation of the α/β-tubulin heterodimer. Using molecular dynamics simulations, we examine phosphorylated α-tubulin (α:Ser165-P) and phosphorylated β-tubulin (β:Ser172-P) individually for their structural effects on local and global MT structure. Findings suggest that most structural effects are due to α-tubulin phosphorylation, including increased H-bonding that engenders a closer, more stable interaction between the γ-phosphate of β:GTP and nearby residues in β:tubulin. Consequently, these increased interactions lead to impaired accessibility of the GTP γ-phosphate to hydrolysis.

## RESULTS

Effects of phosphorylation of α:Ser165 or β:Ser172 were investigated with the 6-mer model (**Figure 2A**), which consists of three pairs of heterodimeric chains (AD, CF, BE) with chain CF standing for the central dimer. We are specifically interested in the structural changes in the inter-dimer cleft (the E-site) upon phosphorylation. Note that in what follows, only the results of the central dimer (chain CF) are presented, primarily because it is more representative of the heterodimers embedded in the infinitely large MT structures. Furthermore, quantitative analyses showed that the inclusion of the dimers on the edge (chains AD and BE) suppresses the observed impacts of phosphorylation because of their elevated hydration compared to the central CF dimer (**Figure S2** in the “Supporting Information”).

**Figure 2.**
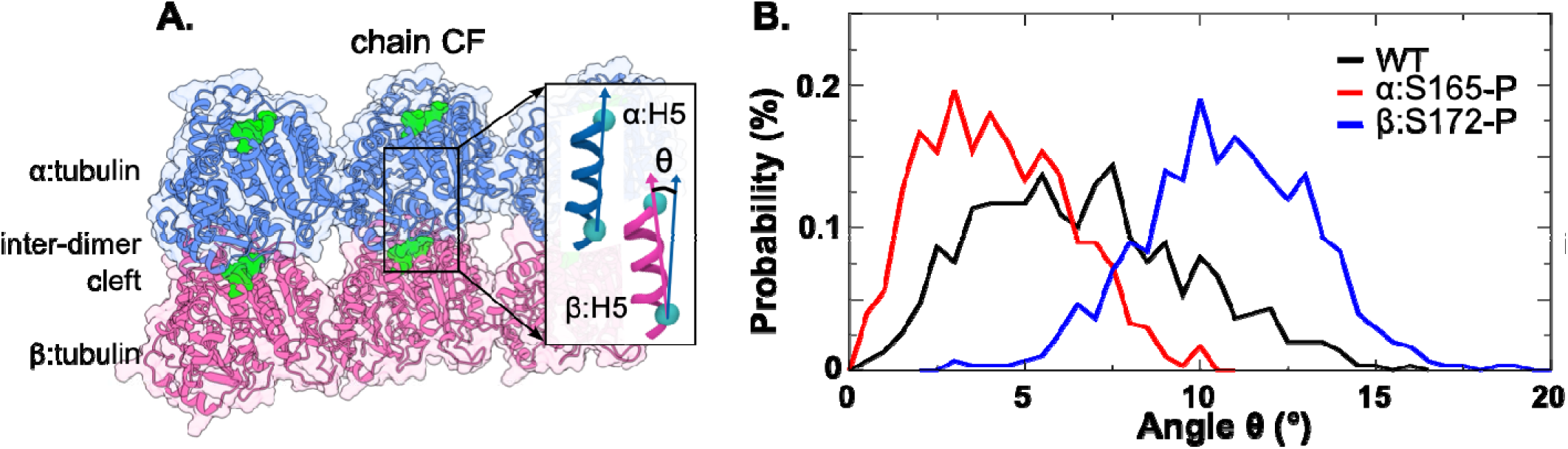
A) Representation of the 6-mer heterodimer system in which α-tubulin and β-tubulin are shown in blue and magenta, respectively. Chain CF stands for the central heterodimer. B) The bending angle (θ) at the inter-dimer cleft of each CF chain for all variants, which is defined using the H5 helices on the α- and β-tubulins demonstrated in the inset of panel (A).

### Phosphorylation impacts MT bending

MT structure is stabilized by protein-protein interactions arising from lateral interactions between homodimers, and longitudinal contacts between heterodimers, both depending on the state of the E-site nucleotide (β:GTP *vs*. β:GDP). The structure of polymerizing microtubules with β:GTP has characteristically straight protofilaments, while microtubules undergoing depolymerization are in the β:GDP state and exhibit a curved appearance (6).

Evaluation of MT structure for each phosphorylated state was performed by measuring the angle defined at the inter-dimer cleft (**Figure 2B**). Specifically, the angle represents the inter-subunit orientation that is obtained with vectors defined by the Cα atoms of two amino acids in the α:H5 helix (Val182 and Thr193) and the corresponding Cα atoms in the β:H5 helix (Val182 and Gln193), which are roughly parallel in an α/β- tubulin dimer (6). Compared with the WT system, which displayed a dominant angle of 7.5°, the ⍰:Ser165-P system exhibited an angle of 3.0°, indicating a more extended protofilament, whereas the β:Ser172-P gave a more curved angle of 10°, which is consistent with a depolymerizing protofilament (6). These findings imply that the global MT architecture can be regulated by phosphorylation that favors either an extended (α:Ser165-P) or a bent conformational state (β:Ser172-P).

### Phosphorylation causes displacement of secondary structures at the inter-dimer cleft

Phosphorylation of α-tubulin increases the distance between the phosphorylated site at α:Ser165-P and the α:H8 helix (α:Ser165-P:Cα - α:Glu254:Cα and α:Ser165-P:Cα - α:Gln256:Cα) (**Table 1**). This movement brings the residues in α:H8 helix significantly closer to the γ-phosphate (Pγ) of β:GTP. Specifically, the Cα in α:Glu254 is 1.1 Å closer and α:Gln256 is 1.6 Å closer in the α:Ser165-P system compared to those in WT (**Table 1**). Phosphorylation of α:Ser165 also increases the distance between the carboxylate oxygens of α:Glu254 (OE1/OE2) and the γ−phosphate of β:GTP, giving a value of 6.6 Å for α:Ser165-P *vs* 5.7 Å for WT (**Table 1**). That is, compared with WT, the phosphorylation of α-tubulin brings the backbone of the α:H8 helix closer to the γ-phosphate of β:GTP, and meanwhile repels the carboxylate oxygens of the catalytic residue α:Glu254. Consequently, the side chain of α:Glu254 exhibits a different (i.e., relatively distorted) orientation towards β:GTP in the α:Ser165-P system. On further inspection of the local structure suggests a vertical orientation of this carboxylate group relative to the γ-phosphate in β:GTP in the α:Ser165-P system (**Figure 3A)** which could disfavor hydrolysis of the γ-phosphate of β:GTP.

**Figure 3.**
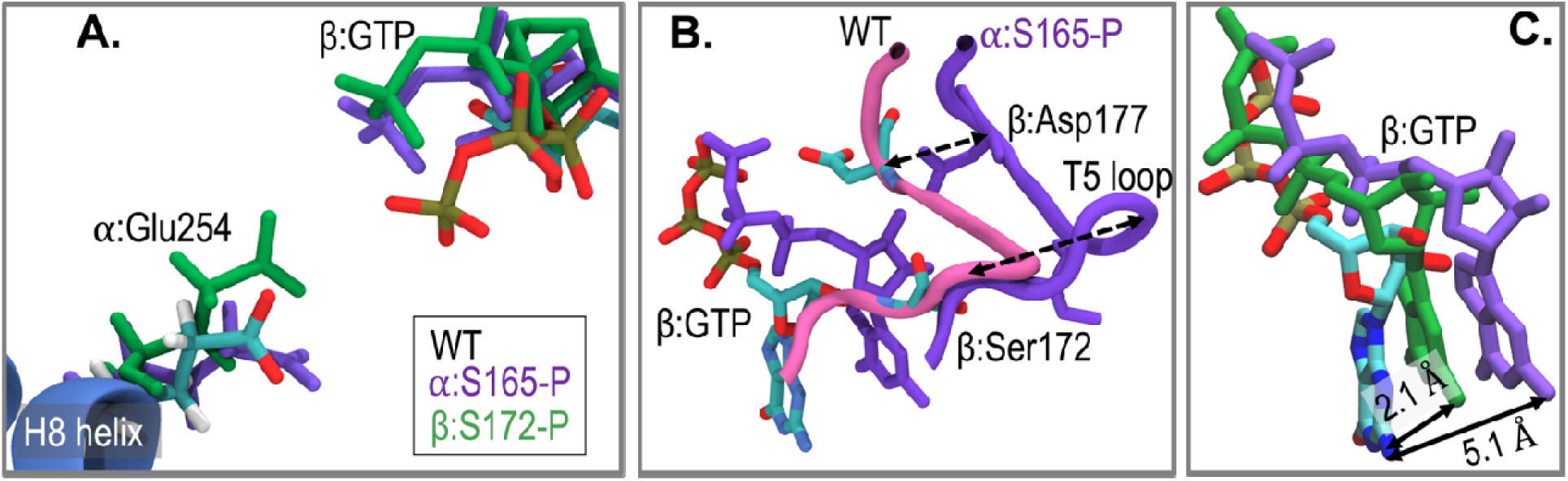
Conformational changes induced by phosphorylation at the inter-dimer cleft. A) The carboxylate group (COO^-^) is roughly vertically oriented towards the γ-phosphate of β:GTP in α:Ser165-P, whereas one carboxylate oxygen is oriented towards the γ-phosphate atom in the WT and β:Ser172-P systems. B) Structural impact of phosphorylated α-tubulin (α:Ser165-P) on the position of β:GTP and loop T5 (β:Asp177). C) Overlay of the structures of β:GTP that illustrates its altered alignment, especially in the α:Ser165-P system. All superimposed structures were aligned using the backbone of helix H8 of α-tubulin.

**Table 1:**
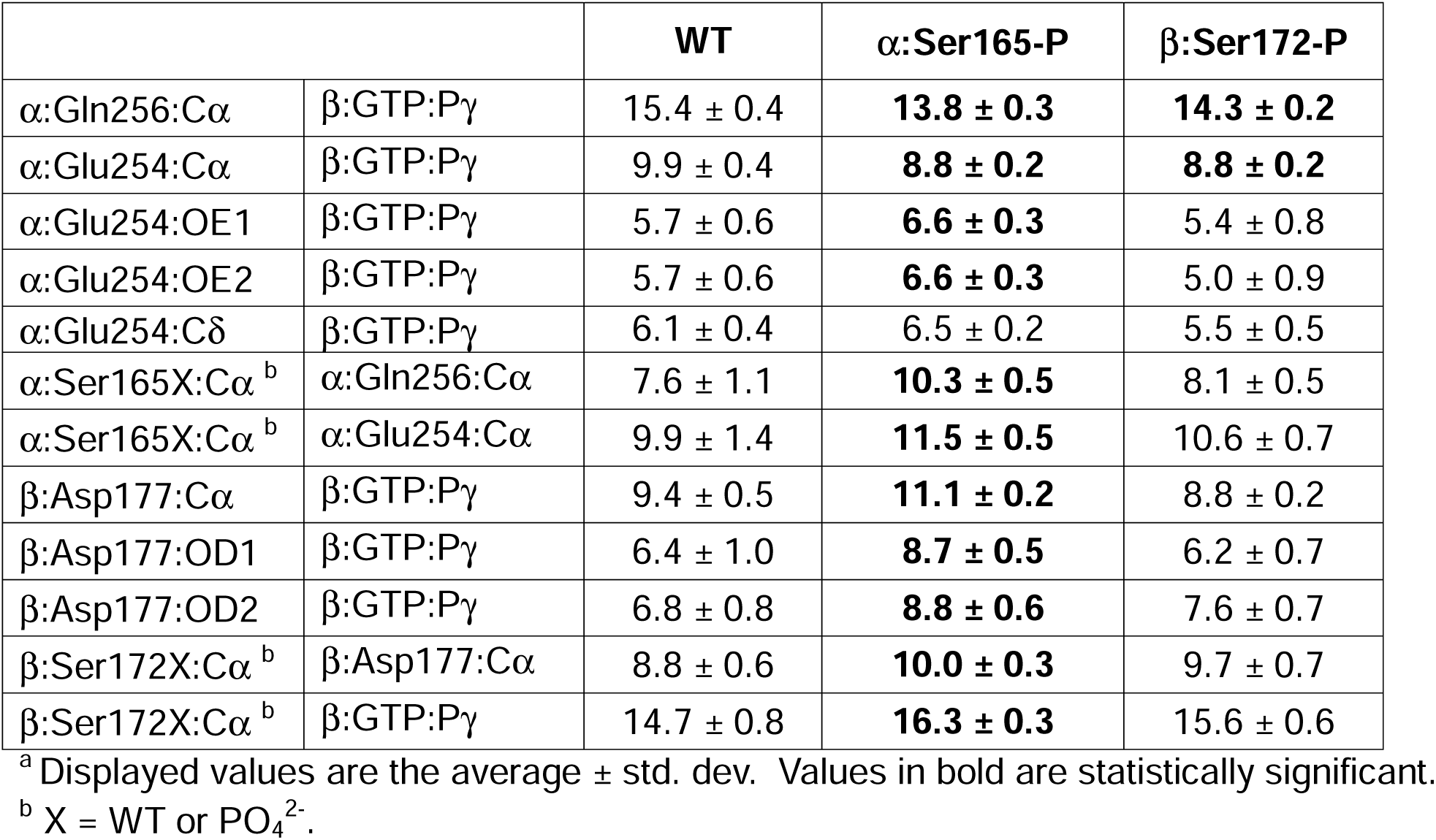
Distances (Å) between residues of the central dimer of the WT, α:Ser165-P, and β:Ser172-P variants with bound β:GTP. ^a^.

| | | WT | $\alpha$:Ser165-P | $\beta$:Ser172-P |
| --- | --- | --- | --- | --- |
| $\alpha$:Gln256:Cα | $\beta$:GTP:Pγ | 15.4 ± 0.4 | **13.8 ± 0.3** | **14.3 ± 0.2** |
| $\alpha$:Glu254:Cα | $\beta$:GTP:Pγ | 9.9 ± 0.4 | **8.8 ± 0.2** | **8.8 ± 0.2** |
| $\alpha$:Glu254:OE1 | $\beta$:GTP:Pγ | 5.7 ± 0.6 | **6.6 ± 0.3** | 5.4 ± 0.8 |
| $\alpha$:Glu254:OE2 | $\beta$:GTP:Pγ | 5.7 ± 0.6 | **6.6 ± 0.3** | 5.0 ± 0.9 |
| $\alpha$:Glu254:Cδ | $\beta$:GTP:Pγ | 6.1 ± 0.4 | 6.5 ± 0.2 | 5.5 ± 0.5 |
| $\alpha$:Ser165X:Cα <sup>b</sup> | $\alpha$:Gln256:Cα | 7.6 ± 1.1 | **10.3 ± 0.5** | 8.1 ± 0.5 |
| $\alpha$:Ser165X:Cα <sup>b</sup> | $\alpha$:Glu254:Cα | 9.9 ± 1.4 | **11.5 ± 0.5** | 10.6 ± 0.7 |
| $\beta$:Asp177:Cα | $\beta$:GTP:Pγ | 9.4 ± 0.5 | **11.1 ± 0.2** | 8.8 ± 0.2 |
| $\beta$:Asp177:OD1 | $\beta$:GTP:Pγ | 6.4 ± 1.0 | **8.7 ± 0.5** | 6.2 ± 0.7 |
| $\beta$:Asp177:OD2 | $\beta$:GTP:Pγ | 6.8 ± 0.8 | **8.8 ± 0.6** | 7.6 ± 0.7 |
| $\beta$:Ser172X:Cα <sup>b</sup> | $\beta$:Asp177:Cα | 8.8 ± 0.6 | **10.0 ± 0.3** | 9.7 ± 0.7 |
| $\beta$:Ser172X:Cα <sup>b</sup> | $\beta$:GTP:Pγ | 14.7 ± 0.8 | **16.3 ± 0.3** | 15.6 ± 0.6 |

<sup>a</sup> Displayed values are the average ± std. dev. Values in bold are statistically significant.
<sup>b</sup> X = WT or PO<sub>4</sub><sup>2-</sup>.

Phosphorylation of α:Ser165 also affects β:loop T5 (aa 171-183), a flexible structural element that participates in longitudinal interactions between heterodimers (**Figure 3B**) and overlaps with the target site (aa 175-213) for MT-destabilizing anti-cancer drugs such as vinblastine (28). Phosphorylated α:Ser165 moves β:Asp177 (loop T5) away from the γ−phosphate of β:GTP by 1.7 Å [*i.e*., moving its position from 9.4 Å (WT) to 11.1 Å (α:Ser165-P)]. (**Table 1** and **Figure 3B**). In contrast, phosphorylated β:Ser172 has a much weaker effect on the position of β:Asp177, which consists of moving it somewhat closer (8.8 Å for β:Ser172-P *vs* 9.4 Å for WT). The finding that α:Ser165-P displaces both β:loop T5 and α:helix H8 is consistent with a previous proposal (12) that helix H8/loop T7 of α-tubulin and the T3/T5 loops in β-tubulin move together as a single structural unit during polymerization, presumably to strengthen longitudinal interactions at the inter-dimer cleft.

### Phosphorylation of α-tubulin shifts the position of the β:GTP nucleotide and strengthens its H-bonding to β-tubulin

Phosphorylation of α:Ser165 is predicted to alter the spatial orientation of β:GTP with respect to α:Glu254 (13). To assess this possible change in the phosphorylated systems relative to the WT, each structure was superimposed with respect to its helix H8 in α-tubulin. α:Helix H8 is a useful reference point because it is structurally stable, adjacent to β:GTP, and contains the essential catalytic residue α:Glu254. Relative to the WT, phosphorylation leads to different structural shifts of the nucleotide base of β:GTP from 2.1 Å for β:Ser172-P to 5.1 Å for α:Ser165-P, supporting the stronger impact of α-tubulin phosphorylation on the β:GTP nucleotide (**Figure 3C**). We calculated several representative distances (**Table 1**), all of which support the stronger influence of α-tubulin phosphorylation compared to β-tubulin phosphorylation.

In describing molecular structures, hydrogen bonds (HBs) are essential in terms of local steric and structural effects, molecular stability, and their effects on biological function (29–32). A consideration of the average number of hydrogen bonds formed between β-tubulin residues and the phosphate oxygen atoms of β:GTP reveals an altered pattern of H-bonding network to β:GTP before and after phosphorylation (**Table 2)**, most notably in the α:Ser165-P system. In this system, a significantly higher number of hydrogen bonds occur that involve the γ-phosphate oxygens (OG: 3.9 ± 0.8) as compared to those with WT (0 ± 0) and β:Ser172-P (0.4 ± 0.5). See **Figure S3** in the “Supporting Information” for the naming rule of the GTP atoms. Moreover, the numbers of hydrogen bonds involving the β-phosphate oxygens (OB) show a similar trend: a stronger H-bond exists in the α:Ser165-P system, though the differences between the three variants are relatively weaker compared to those involving the γ-phosphate oxygens. The finding of increased H-bonding to β:GTP in the α:Ser165-P system suggests a diminished flexibility of the γ-phosphate group that could suppress its hydrolysis.

**Table 2.**
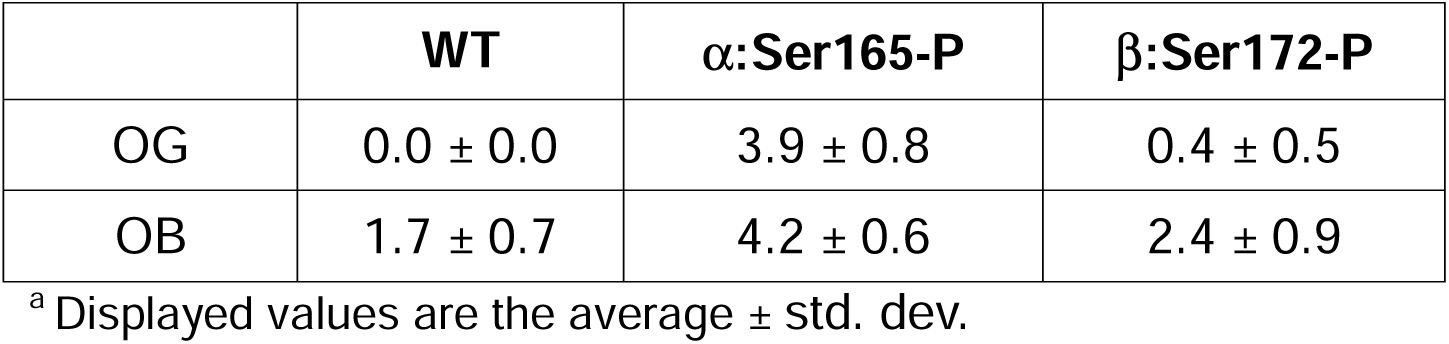
Total number of hydrogen bonds formed between β-tubulin residues and the β-and γ-phosphate oxygen atoms (OG and OB) of β:GTP.^a^.

In addition to the number of H-bonds in **Table 2**, the H-bonding pattern contributed by residues that define the β:GTP-binding pocket undergoes remodeling as a result of phosphorylation (See **Table S1** in the “Supporting Information”.) This was found to apply in response to phosphorylation of α-tubulin (**Table 3**). As a result, the distances between these residues and the γ-oxygens (OGs) and, to a lesser extent, β-oxygens (OBs), were shortened, thus drawing β:GTP deeper into the GTP binding pocket of the α:Ser165-P system.

**Table 3.** Distance (Å) between β:GTP atoms and β-tubulin residues in the β:GTP-binding pocket.

| | | WT | $\alpha$:Ser165-P | $\beta$:Ser172-P |
| --- | --- | --- | --- | --- |
| GTP-OB1 | Ser138-OG | 6.5 ± 0.6 | **2.7 ± 0.2** | 5.9 ± 0.6 |
| GTP-OG1 | Thr143-N | 5.4 ± 0.1 | **3.0 ± 0.2** | **3.4 ± 0.5** |
| GTP-OG1 | Thr143-OG1 | 6.2 ± 0.2 | **2.7 ± 0.1** | 4.2 ± 0.7 |
| GTP-OG3 | Asn99-N | 7.2 ± 0.1 | 3.9 ± 0.4 | 5.0 ± 0.1 |
| GTP-OG3 | Asn99-ND2 | 6.2 ± 0.2 | **2.8 ± 0.2** | 3.7 ± 1.0 |
| GTP-OG3 | Gly142-N | 4.8 ± 0.7 | **3.4 ± 0.3** | 3.5 ± 0.3 |

<sup>a</sup> Displayed values are the average ± std. dev. Values in bold represent distances that are < 3.5 Å and therefore likely to form an H-bond. See **Figure S3** in the Supplemental Information for the naming rules.

The analysis of interaction energies between H-bond donors and β:GTP (**Table 4**) reinforces these observations by displaying varying strengths of interactions on the inter-dimer cleft. α:Ser165-P exhibits the most stable (most negative) overall interactions between β:GTP and the β-tubulin residues that form H-bonds between them.

**Table 4.**
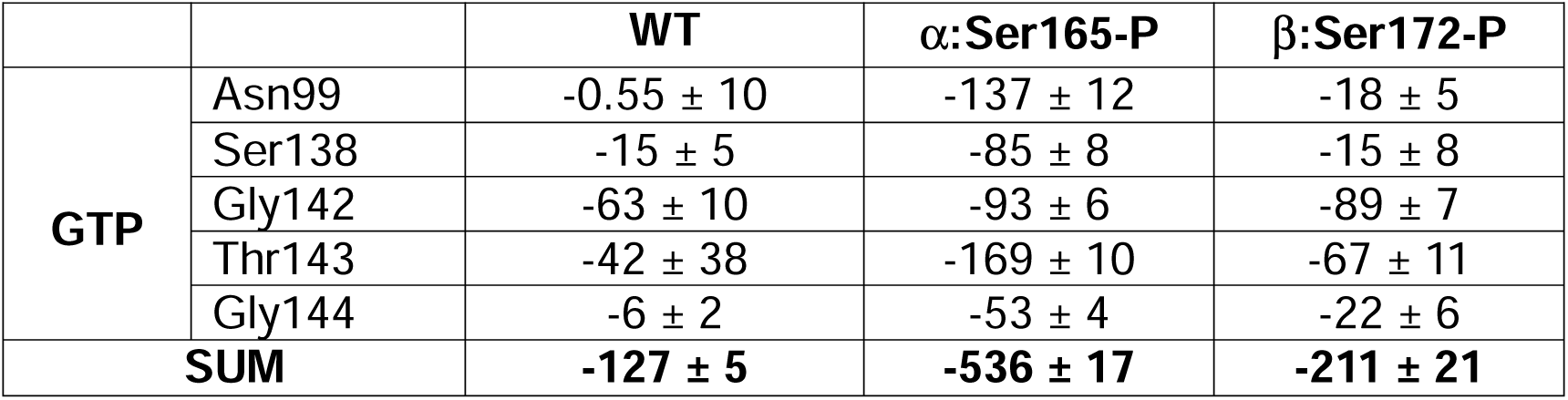
Interaction energies (kJ/mol) between β:GTP and selected β-tubulin residues that form H-bonds to β:GTP.

|  |  | **WT** | **$\alpha$:Ser165-P** | **$\beta$:Ser172-P** |
| --- | --- | --- | --- | --- |
| **GTP** | Asn99 | -0.55 ± 10 | -137 ± 12 | -18 ± 5 |
| | Ser138 | -15 ± 5 | -85 ± 8 | -15 ± 8 |
| | Gly142 | -63 ± 10 | -93 ± 6 | -89 ± 7 |
| | Thr143 | -42 ± 38 | -169 ± 10 | -67 ± 11 |
| | Gly144 | -6 ± 2 | -53 ± 4 | -22 ± 6 |
| **SUM** |  | **-127 ± 5** | **-536 ± 17** | **-211 ± 21** |

These results support a model in which α:Ser165-P promotes remarkably stronger β:GTP binding by H-bonding with these residues, while β:Ser172-P only marginally enhances such β-tubulin-β:GTP interactions. Therefore, phosphorylation of α-tubulin generates the strongest H-binding energy to β:GTP through interactions with neighboring residues.

These results are consistent with the stabilities of β:GTP measured by RMSD values in the WT, α:Ser165-P, and β:Ser172-P systems, as illustrated in **Figure 4**. Each RMSD analysis was performed with reference to the Cα atoms of all residues in β-tubulin that define the β:GTP-binding pocket. During the 1.2-1.6 μs period, the average RMSD value identified for each system showed that the α:Ser165-P system exhibits the lowest average fluctuation (0.10 ± 0.01 nm, highest stability) when compared to the β:Ser172-P (0.13 ± 0.02 nm) and WT (0.18 ± 0.04 nm) systems. Thus, in the α:S165-P system, the increased H-bonding (number and energy in **Tables 2-4**) between β:GTP and its binding pocket leads to a suppressed fluctuation of the β:GTP in the α:Ser165-P system. In contrast, in the β:Ser172-P and WT systems, β:GTP has a less constraining H-bonding environment and is likely to be more flexible.

**Figure 4.**
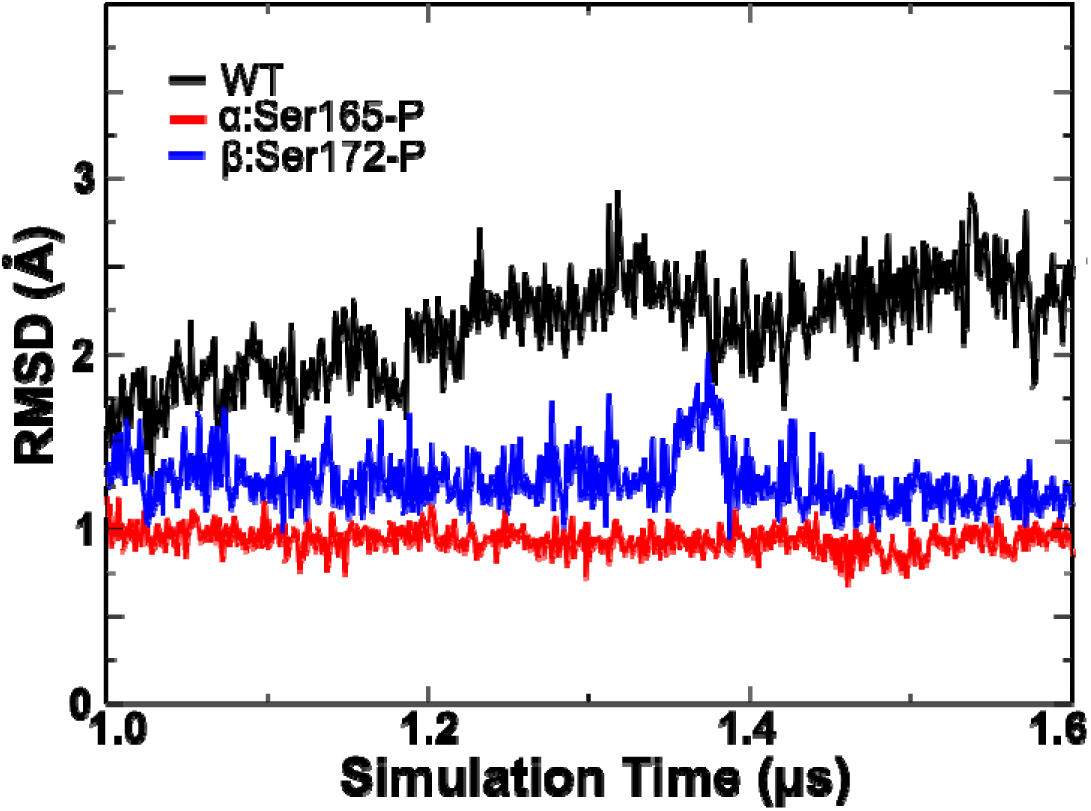
RMSD plots describe fluctuations of β:GTP in the WT, α:Ser165-P, and β:Ser172-P systems. The α:Ser165-P system exhibits the highest stability with the lowest average RMSD value of 0.10 ± 0.01 nm, compared with β:Ser172-P with a value of 0.13 ± 0.02 nm, and WT with a value of 0.18 ± 0.04 nm, being the least stable system.

We further calculated the covariance of the interfacial residues in the E-site (**Figure S4** in the “Supporting Information”). A weaker correlation between the α:H8 helix and the β:tubulin residues in the α:Ser165-P system is observed, compared with the WT and β:Ser172-P systems. Again, this is consistent with the increased distance between the α:H8 helix residues α:Glu254:OE1/OE2 and β:GTP (Τ**able 1**) and stronger binding between β:GTP and β:tubulin (**Tables 2-4**).

### Phosphorylation of α /β-tubulin and GTP hydrolysis

A previous report (13) experimentally demonstrated by an immunochemical assay that phosphorylated α-tubulin increases the signals related to β:GTP in β-tubulin, as compared with WT. This result is consistent with phosphorylated α-tubulin leading to suppressed β:GTP hydrolysis. Hydrolysis of the γ-phosphate of β:GTP is expected to occur via a network of water molecules that forms between α:Glu254 and β:GTP within the inter-dimer cleft. First, the average distance between the carboxylate carbon atom (Cδ) of α:Glu254 and the Pγ atom of β:GTP was calculated for each system. These averages were: 6.1 ± 0.4 Å in the WT system, 6.5 ± 0.2 Å in the α:Ser165-P system, and 5.5 ± 0.5 Å in the β:Ser172-P system (**Table 1**). Representative snapshots of each system (**Figure 5**) were selected to view interacting water molecules between α:Glu254 and the γ-phosphate of β:GTP. A water-mediated pathway is supposed to promote the hydrolysis reaction of the γ-phosphate via the carboxylate group of α:Glu254. In the α:Ser165-P system, we noted the vertical orientation between the carboxylate group of α:Glu254 and the γ-phosphate of β:GTP, as well as a somewhat longer distance between them. Both features are accompanied by fewer water-mediated pathways connecting them and consequently are expected to disfavor β:GTP hydrolysis. In contrast, two water-mediated H-bond-assisted pathways are observed in the β:Ser172-P system, and one in the WT system.

**Figure 5.**
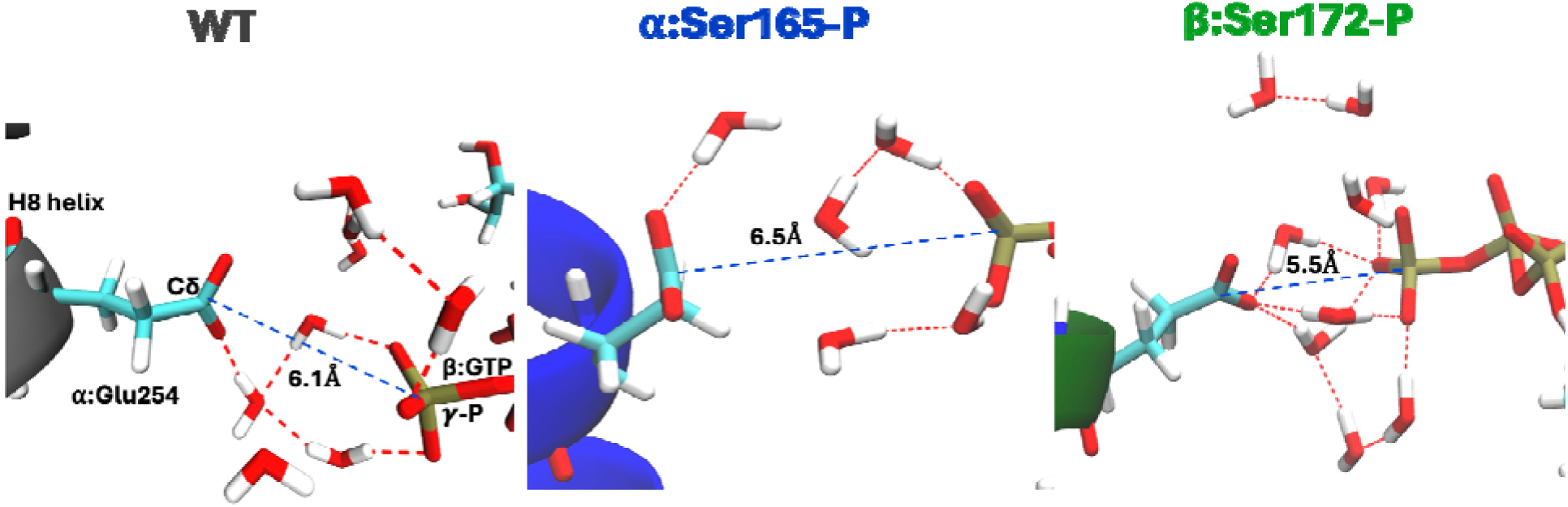
Representative structures of water-mediated network between the carboxylate of α:Glu254 and the γ-phosphate of β:GTP. Each system is defined by the average distance (in Å) between α:Glu254:Cδ and γ-phosphate of β:GTP. Red dashed lines represent hydrogen bonds, while the blue dashed line indicates the distance between α:Glu254:Cδ and the γ-phosphate of β:GTP. β:Ser172-P has an extensive water-mediated network with two observable paths that link α:Glu254 to β:GTP, whereas α:Ser165-P displays no such paths. Additional snapshots are shown in **Figure S5** in the “Supporting Information”.

## DISCUSSION

Previous efforts demonstrated the effects on cellular activity that are associated with PKC-mediated phosphorylation of Ser165 in α-tubulin (9, 10, 13), or CDK1-mediated phosphorylation of Ser172 in β-tubulin (14, 33). In-depth analysis of the impact of α-tubulin phosphorylation on MT dynamics by live cell imaging supported its role in promoting an increased rate and duration of polymerization (10). This effect correlated with elevated cell motility, an aspect of metastatic potential. However, phosphorylation of β-tubulin by CDK-1 promotes cell proliferation. The extent to which PKC and CDK-1 compete with each other for their sites of phosphorylation (as in a toggle switch) was made evident by a comparison of blocked α-tubulin phosphorylation (with α:S165N), which suppressed motility of metastatic breast cells while causing hyper-proliferation in an animal model, presumably due to phosphorylated β-tubulin (9,11). To explain these effects, the present study explored a structural basis for how phosphorylation of each site could result in the expected impact on MT dynamics.

Hallmark structural changes identified *in silico* for phosphorylated α-tubulin include 1) linearization of the microtubule structure that is consistent with a polymerizing MT, 2) displacement of secondary structures within α-tubulin and β-tubulin (α:H8 helix, β:T5 loop), 3) shift in the nucleotide moiety of β:GTP, 4) immobilization of β:GTP by increased H-bonding to the γ-phosphate, and 5) decreased capacity for β:GTP hydrolysis. A prominent mechanistic feature of α:Ser165-P is that these events affect the γ-phosphate of β:GTP. The following model is proposed: the higher number and stabilizing interactions of H-bonds that bind β:GTP (**Tables 2-4**) embed it deeper into the β:GTP-binding pocket, as well as pull it away from the inter-dimer cleft. In turn, the increased distance between α:Glu254:Cδ and the γ-phosphate interrupts the water network. Based on the known phenotypic consequences on microtubule dynamics by α:Ser165 phosphorylation in living cells (10), the identified computational features are consistent with decreased β:GTP hydrolysis (13) and a prolonged phase of MT polymerization (10).

Conversely, in β-tubulin, it was shown that β:Ser172-P binds β:GTP more loosely (**Tables 2-4**), similar to a previous report (14), and the increased curvature of its global structure (**Figure 2**) is consistent with a depolymerizing microtubule. Furthermore, the multiple opportunities afforded by the water network to perform β:GTP hydrolysis in β:Ser172-P (**Figure 5**) lend strong support for this model. Taken together, the foregoing computer-based analyses identify key structural details that are consistent with known functional characteristics of phosphorylated α- or β-tubulin and their associated effects on MT function.

This computational study sought to formalize the structural responses to phosphorylation of α:Ser165 or β:Ser172 at the inter-dimer cleft. The identified elements that operate within α- or β-tubulin at this site and β:GTP provide new insights into how dynamic instability may be modulated. As a practical application, these structural details may be helpful in screening mutations in tumor genomic databases for similar perturbations and thus provide a tool to predict metastatic or tumorigenic potential.

We propose that α/β-tubulin provides a hub for the activities of PKC and CDK1 and perhaps other protein kinases. An attractive model is that phosphorylation of either α-tubulin (by PKC) or β-tubulin (by CDK1) is a response to the prevailing protein kinase activity. The toggle switch hypothesis thus offers a framework for understanding how intracellular signaling pathways regulate MT dynamic instability and consequently specific cellular phenotypes.

## EXPERIMENTAL PROCEDURES

### Tubulin Variants Investigated

The wild-type (WT) system with β:GTP was built from a cryo-EM structure of a GMPCPP-stabilized microtubule in the presence of Taxol at 4.7 Å resolution (PDB ID: 3J6E) (12). Non-hydrolyzable GTP analogue GMPCPP (G2P) was converted to GTP by replacing C3A with O3A (13). AlphaFold2 (15) was employed to fill in missing residues in the cryo-EM structure. AlphaFold2 is a deep learning model that utilizes evolutionary information to predict protein structure and missing sequences with high accuracy (16). For 3J6E, AlphaFold2 was used to fill in the missing residues 39 to 48 on the α-tubulin. From the resulting structure, three α/β heterodimers were extracted to construct the WT 6-mer system.

It is noted here that in 3J6E.pdb, the numbering of β-tubulin residues is shifted higher by 2 residues so that Ser172 occurs as Ser174. To conform with the literature, the numbering of β-tubulin residues has been adjusted to standard values. Phosphorylated variants (α:Ser165-P and β:Ser172-P) were prepared using the open-source programs PyMOL, VMD, and in-house scripts.

### The 6-mer System

To examine the different states of MT assembly, we simulated a “6-mer” system for each variant, representing the central region of MT protofilaments (PFs) (**Figure 1**). Each 6-mer consists of fragments of three protofilaments, each with one α/β heterodimer and infinitely repeating across the Z-dimension of the simulation box. The system was dissolved in aqueous solution in a three-dimensional periodic boundary constraint (PBC) box using the recommended CHARMM TIP3P water model with water structures constrained via the SETTLE algorithm (17) and 0.20 M KCl solution, as conducted previously (13). K^+^ ions were also added as counterions to neutralize the whole system.

### Atomistic Molecular Dynamics Simulations

All-atom explicit solvent molecular dynamics (AA-MD) simulations were carried out using the software GROMACS (version 2024.5) (18). The CHARMM 36m all-atom force field (19) was employed, which has been extensively employed for protein simulations (20–23) and provides the force field parameters for all components employed here: proteins, GTP, GDP, and metal ions.

The energy of the system was first minimized using the steepest descent algorithm, which was followed by further equilibration using the NVT ensemble (constant number of particles, volume, and temperature) of 10 ps of simulation with the timestep of 1 fs, the NPT ensemble (constant number of particles, pressure, and temperature) of 10 ps simulation with the timestep of 1 fs, and 1 ns of NPT simulation using the timestep of 2 fs with all hydrogen-involved covalent bonds constrained through the LINCS algorithm (24, 25). In these equilibrium simulations, position restraints were employed on the non-hydrogen atoms of the proteins and the GTP ligands, with the restraint force constant of 1000 kJ/mol/nm^2^. Such restraints were removed in the production simulation below, allowing their full relaxation.

Each production was run for 1600 ns using a 2 fs timestep. The energies and coordinates were saved every 1 ns. The trajectories from the last 600 ns were used for data analyses. Verlet cutoff schemes were updated every 20^th^ timestep. The short-range Coulombic interactions were computed using the Potential Shift Verlet Modifier up to 1.2 nm, with the long-range interactions calculated using the Particle Mesh Ewald (PME) method (26) with a Fourier spacing of 0.12 nm. The Lennard-Jones interactions were switched off between 1.0 – 1.2 nm using the Potential Switch method.

The temperature coupling was employed using the Nosé-Hoover algorithm with the temperatures of water and non-water molecules coupled separately at 298 K with a time constant of 1 ps. The pressure was managed semi-isotropically using the Parrinello–Rahman barostat with a time coupling constant of 4 ps, compressibility of 4.5 × 10^-5^ bar^1^, and reference pressure of 1 bar (27). The root-mean-square displacement (RMSD) calculations supported that the systems are roughly converged within 400 – 800 ns of the simulation (**Figure S1** in the “Supporting Information”).

## DATA AVAILABILITY

All data are contained within the manuscript.

## SUPPORTING INFORMATION

This article contains supporting information.

## ACKNOWLEDGEMENTS

This work is supported by National Institutes of Health grant GM153696 (to S.A.R.).

## ABBREVIATIONS

αSer165-P: phosphorylated α-tubulin
βSer172-P: phosphorylated β-tubulin
MT: microtubule
GTP: guanosine triphosphate
PKC: protein kinase C
CDK-1: cyclin-dependent kinase-1

