## Supporting Info for "How Phosphorylation of α/β-Tubulin Perturbs Microtubule Structure: A Computational Study"

Annemarie Ianos^1^, Ahmed Osman^2,3^, Baofu Qiao^1,2,*^, and Susan A. Rotenberg^2,3,*^

^1^Department of Natural Sciences, Baruch College, City University of New York, New York, New York 10010, USA.

^2^Graduate Center, City University of New York, New York, New York 10010, USA.

^3^Department of Chemistry & Biochemistry, Queens College, City University of New York, New York, New York 10010, USA.

^*^ Corresponding authors:

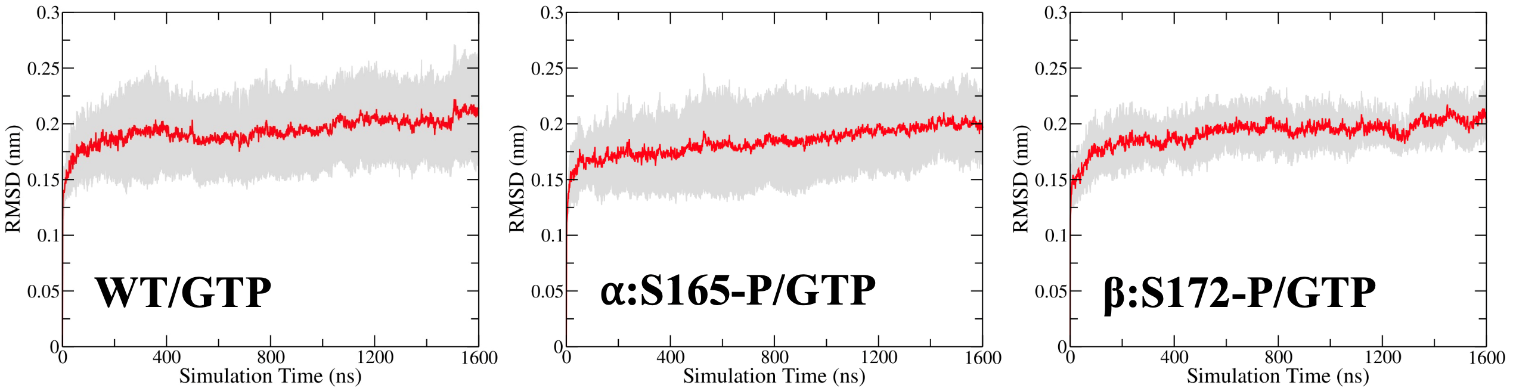

**Figure S1.** Average RMSD across all dimers in the different variants investigated.

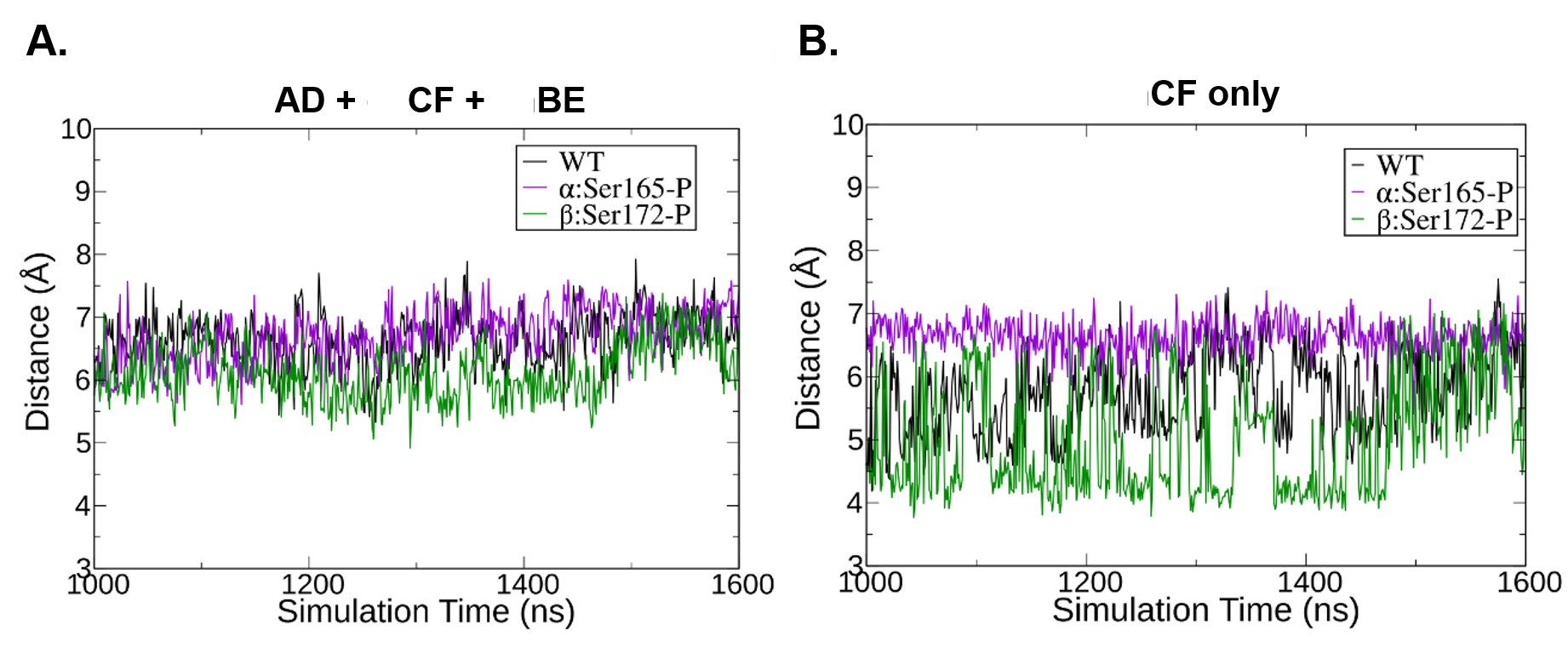
 **Figure S2.** Time dependence of the distance between β:GTP:Pγ and α:Glu254:Cδ. (A) data for all chains; (B) data for the center dimer only (chain CF).

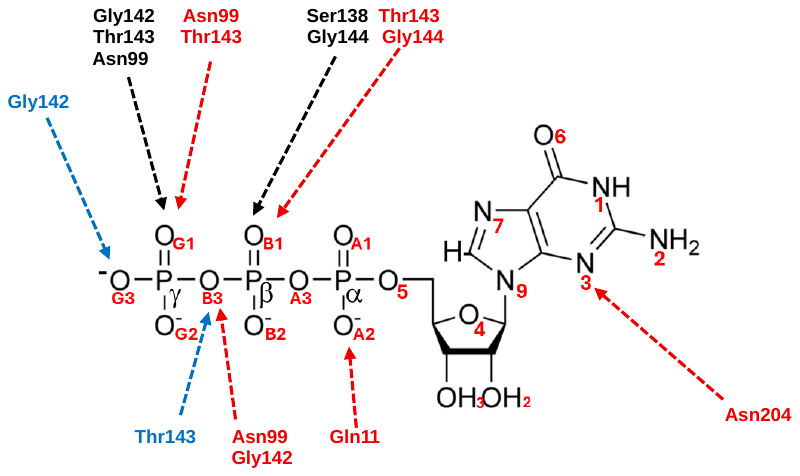

**Figure S3.** Gain/loss of H-bonds in β:GTP due to phosphorylation of α- or β-tubulin. Residues shown in **black** refer to H-bonds formed by residues in the α:Ser165-P system. Residues shown in **red** were eliminated in the β:Ser172-P system, whereas residues shown in **blue** were formed in the β:Ser172-P system.

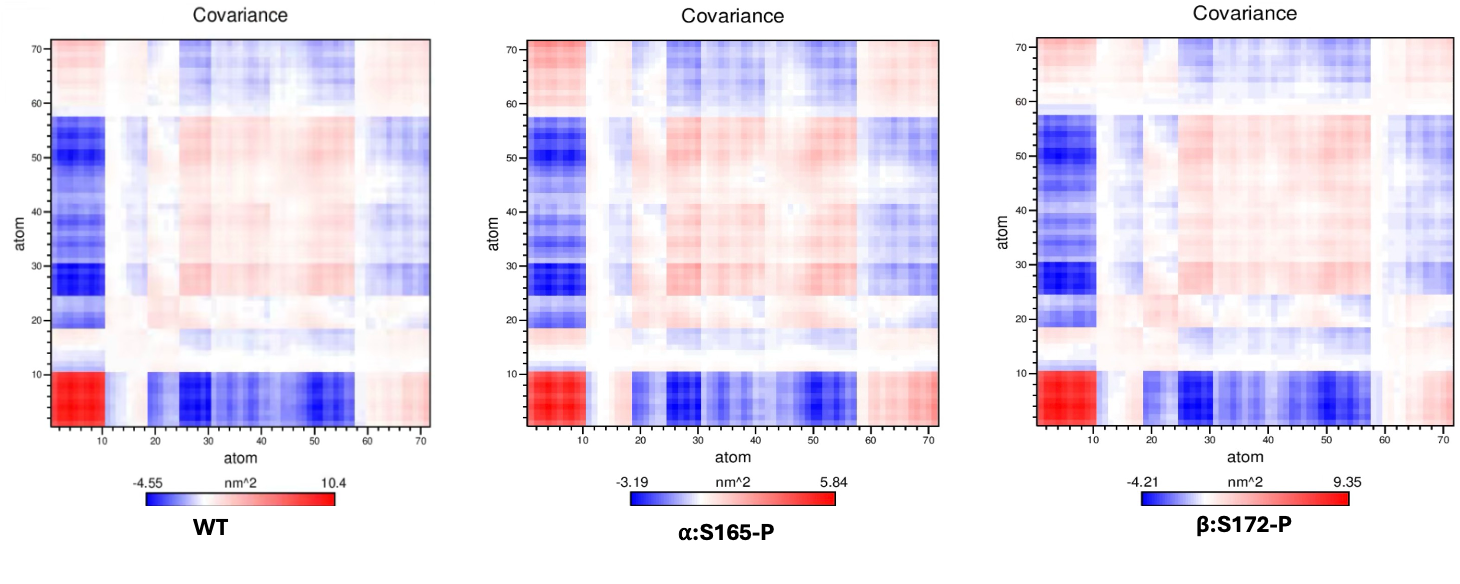

**Figure S4. Covariance of the interfacial residues in WT, α:Ser165-P, and β:Ser172-P.** Atoms 1-10 refer to the Cα-atoms on the α-subunit of residue α:Ser165 and the H8 helix (L252-V260); the other atoms refer to the Cα-atoms on the β-subunit: β:Ser172 and the interfacial residues within 6 Å of β:GTP.

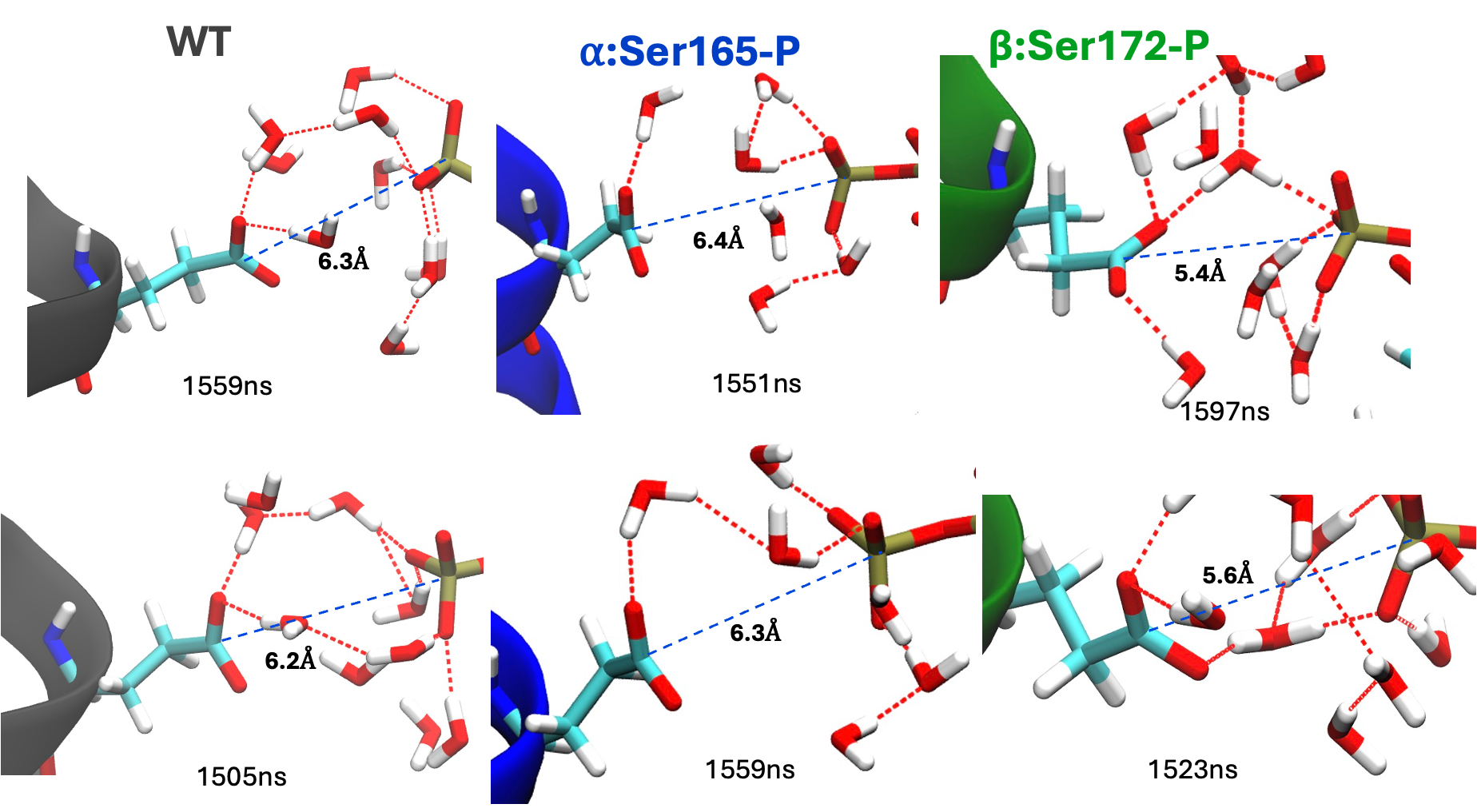

**Figure S5.** Additional structures illustrating the water-mediated interaction network between α:Glu254 and β:GTP. Representative snapshots were selected from the 1000–1600 ns analysis window: WT (1559 and 1505 ns), α:Ser165-P (1551 and 1559 ns), and β:Ser172-P (1597 and 1523 ns). Only in the α:Ser165-P system are α:Glu254 and γ-phosphate of β:GTP oriented vertically, leading to decreased interactions between them. WT and α:Ser165-P exhibit a similar distance between α:Glu254-Cδ and γ-phosphate of β:GTP, whereas in the β:Ser172-P system, α:Glu254 is closer to the γ-phosphate and establishes multiple pathways with water molecules.

**Table S1** Percentage existence of H-bonds between the β:GTP and residues in β−tubulin in the phosphorylated 6-mer mutants and wildtype (WT). Amino acid residues reflect standard numbering.

| **Hydrogen Bond (%)** | **WT** | **α:Ser165-P** | **β:Ser172-P** |
| --- | --- | --- | --- |
| Asn226-OD1 to GTP-N1 | 99.2 | 98.5 | 92.2 |
| Asn204-OD1 to GTP-N2 | 97.0 | 96.1 | 88.0 |
| Asn204-ND2 to GTP-N3 | 68.7 | 63.9 | 42.7 |
| Gly144-N to GTP-OB1 | 66.2 | 89.6 | 38.9 |
| Ser138-OG to GTP-OB1 | 0.5 | 32.3 | 1.6 |
| Thr143-N to GTP-OG1 | 11.7 | 30.3 | 1.0 |
| Thr143-OG1 to GTP-OG1 | 58.1 | 66.6 | 33.9 |
| Asn99-N to GTP-OG3 | 9.9 | 50.0 | 0.0 |
| Asn99-ND2 to GTP-OG3 | 40.8 | 64.8 | 25.6 |
| Gly142-N GTP-OG3 | 1.9 | 42.5 | 16.9 |
| Glu181-OE to GTP-O3 | 19.7 | 18.7 | 25.0 |
| Asn226-ND2 to GTP-O6 | 95.8 | 95.7 | 94.3 |
| Gln11-NE2 to GTP-N7 | 8.1 | 13.3 | 4.2 |
| Gln11-NE2 to GTP-OA2 | 25.2 | 23.3 | 6.3 |
| Gln15-NE2 to GTP-O6 | 5.7 | 3.4 | 12.8 |
| Thr143-OG1 to GTP-OB1 | 9.2 | 0.0 | 0.4 |
| Thr143-OG1 to GTP-OB3 | 0.3 | 0.8 | 27.0 |
